# A novel biomechanical indicator for impaired ankle dorsiflexion function during walking in individuals with chronic stroke

**DOI:** 10.1101/2023.04.28.538758

**Authors:** Shraddha Srivastava, John H Kindred, Bryant A. Seamon, Charalambos C. Charalambous, Andrea D. Boan, Steven A. Kautz, Mark G Bowden

**Affiliations:** Ralph H. Johnson Veteran’s Affairs Health Care System, Charleston, SC, USA; Department of Health Sciences and Research, College of Health Professions, Medical University of South Carolina, Charleston, SC, USA; Division of Physical Therapy, College of Health Professions, Medical University of South Carolina, Charleston, SC, USA; Department of Basic and Clinical Sciences, Medical School, University of Nicosia, Nicosia, Cyprus; Center for Neuroscience and Integrative Brain Research (CENIBRE), Medical School, University of Nicosia, Nicosia, Cyprus; Departments of Pediatrics, Neurology, and Public Health Sciences, College of Medicine, Medical University of South Carolina, Charleston, SC, USA; Department of Research, Brooks Rehabilitation, Jacksonville, FL, USA

**Keywords:** stroke, ankle angular velocity, ankle angular acceleration, dorsiflexors, gait

## Abstract

Ankle dorsiflexion function during swing phase of the gait cycle contributes to foot clearance and plays an important role in walking ability post-stroke. Commonly used biomechanical measures such as foot clearance and ankle joint excursion have limited ability to accurately evaluate dorsiflexor function in stroke gait. We retrospectively evaluated ankle angular velocity and ankle angular acceleration as direct measures for swing phase dorsiflexor function in post-stroke gait of 61 chronic stroke survivors. Our linear regression models revealed that peak ankle angular velocity (AAV_P_), peak ankle angular acceleration (AAA_P_), peak dorsiflexion angle (DFA_P_) and peak foot clearance (FCL_P_) during swing had a significant relationship (p < 0.05) with impaired dorsiflexion function. AAA_P_ and DFA_P_ accounted for the most variance of dorsiflexion function. Additionally, AAV_P_, AAA_P_, FCL_P_ during swing, correlated significantly with all clinical outcome measures of walking ability. DFA_P_ during swing had a positive correlation only with FMA-LE. Post-hoc William’s *t*-tests, used to compare the magnitude of difference between two non-independent correlations, revealed that the correlation between all clinical measures and DFA_P_ were significantly weaker than with AAV_P_ and AAA_P_. We also found that correlation between FMA-LE and FCL_P_ was weaker than with AAV_P_ and AAA_P_. We found an excellent test-retest reliability for both AAV_P_ (ICC = 0.968) and AAA_P_ (ICC = 0.947). These results suggest that DFA_P_ may only be associated with non-task specific isolated dorsiflexion movement, but not during walking. FCL_P_ is associated with dorsiflexion function and walking ability measures but not as strongly as AAV_P_ and AAA_P_ possibly because FCL_P_ is influenced by contribution from hip and knee joint movements during walking. Therefore, we believe that AAV_P_ and AAA_P_ both can be used as reliable measures of impaired dorsiflexion function in post-stroke gait.

## Introduction

Impaired dorsiflexion function is often associated with poor walking ability following stroke. Insufficient dorsiflexion of the paretic ankle is a common post-stroke gait pattern [1] and is associated with asymmetric gait and slower walking speeds [2]. Gait impairments associated with poor dorsiflexion function lead to reduced walking capacity [3] that can result in limited independent community ambulation. Rehabilitation strategies that focus on improving dorsiflexion and adequate ankle movement during swing phase improve walking ability [4-6]. Therefore, measurement and treatment of dorsiflexion function during swing phase of walking is important for gait rehabilitation following stroke.

Despite the significance of dorsiflexion during swing, there is no gold standard quantitative measure to evaluate dorsiflexion function during walking in stroke gait. Dorsiflexion function is measured typically by dorsiflexor strength with a dynamometer or by muscle activity (i.e., amplitude and timing of dorsiflexors) with surface electromyography. Unfortunately, dynamometer measurements cannot be performed during walking and thus do not quantify dorsiflexion function during walking-specific tasks. Surface electromyography (sEMG) activity of dorsiflexors can be measured during walking, but it may not reflect its relationship with dorsiflexion function accurately because swing phase dorsiflexor activity can be influenced by altered plantarflexor activity patterns [7]. The use of sEMG is also limited in rehabilitation clinics due to the lack of adequate time, education, and resources available to clinicians [8]. During a gait cycle, muscles contribute toward producing certain phase-specific biomechanical functions required for a normal walking pattern [9, 10], and altered muscle activity results in deviations from normal biomechanical functions [11, 12]. Evaluating biomechanical outputs as a consequence of dorsiflexor activity may offer improved validity as a measure for dorsiflexor function during walking.

Currently published biomechanical measures used to capture dorsiflexor function during swing include foot clearance and ankle joint excursion, but these measures have limitations when measuring dorsiflexor function in stroke gait. Dorsiflexor activity during swing contributes to foot clearance and reduced dorsiflexion function can result in limited foot clearance post-stroke. However, compensatory mechanisms such as circumduction of the paretic leg or hiking of the pelvis may increase foot clearance [13] without much contribution from dorsiflexors. Therefore, foot clearance can be improved through compensatory mechanisms and may not accurately measure dorsiflexor function. Ankle joint angle excursion profile is often used to evaluate stroke survivors’ walking ability [14, 15]; however, previous literature suggests that angular excursion has a limited ability to accurately discriminate the impaired movement patterns in individuals with neurological disorders due to soft tissue contractures [16], which often limit the total joint range of motion. There is clearly a need for a biomechanical measure that can address the aforementioned concerns and accurately reflect the function of dorsiflexors during walking in individuals with impaired gait patterns.

Kinetic measures during gait offer the ability to measure the mechanical effect of dorsiflexors during the gait cycle. Muscle activity contributes to the net joint power, and ankle joint power can be estimated to evaluate dorsiflexor activity during swing and stance phases of walking. Walking requires that each leg muscle produces an appropriate amount of force to contribute towards joint power responsible for the execution of phase-specific biomechanical functions [17]. Previous literature has demonstrated that inappropriate force generated by plantar flexors during stance phase is associated with walking impairment in stroke survivors [18]. However, during the swing phase dorsiflexor ankle power is very low and difficult to measure clinically, such that it does not likely provide a useful measure of dorsiflexor function. Ankle angular velocity (AAV) and acceleration (AAA), however, are direct mechanical consequences of dorsiflexor activity during walking [19] and can thus be used as a measure of dorsiflexor function during swing. Specifically, the torque generated by dorsiflexors contributes toward net ankle joint power. Net ankle joint power is the product of ankle joint angular velocity and ankle joint torque:

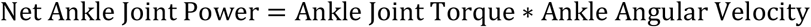

Since joint torque is negligible during swing phase, net ankle joint power is also negligible despite AAV being more substantial and easier to measure directly. This suggests that swing phase ankle angular velocity may serve as a useful measure of dorsiflexor function. Since forces generated by muscles during walking are major contributors towards the acceleration of body segments [9, 19] ankle angular acceleration can be used to evaluate dorsiflexor function during walking in stroke survivors and has the added advantage of being potentially computed using inertial measurement units (IMUs) when inverse dynamic analyses are not available.

The purpose of this study was to investigate whether AAV and AAA are reliable measures to evaluate impaired dorsiflexion function during walking in individuals with chronic stroke. We also evaluated the relationships of paretic AAV and AAA with stroke survivors’ walking ability and determined test-retest reliability of AAV and AAA. We hypothesized that both AAV and AAA will have stronger correlations compared to the commonly used measures of foot clearance and ankle dorsiflexion excursion. Furthermore, we hypothesized that both measures of paretic ankle function will demonstrate acceptable reliability for a clinical measurement tool (ICC > 0.90) [20]. Lastly, we present recommendations for future studies based on the results and the ease of translation into clinical environments.

## Methods

Data from 61 chronic (>6 months) stroke survivors were retrospectively analyzed for this study. Stroke survivors were included if they had residual paresis in the lower extremity (Fugl-Meyer LE motor score ≤34), the ability to walk at least 10 ft. at a self-selected speed ≤1.0m/s and were between the ages of 18-85. Demographic characteristics are reported in Table 1. Since the database used for this retrospective analysis includes data from multiple research studies, we have 1 individual that participated in two different studies at 21 months and 91 months post-stroke respectively. We treated their data as two separate records because their motor and biomechanical responses captured during the first and the second time point were considerably different.

**Table 1:** Participant Demographics in Regression and Correlation Analysis

| **n=61** |  |
| --- | --- |
| **Age** | 60.6 (11.5) [28-85] |
| **Sex** |  |
| Male | 40 / 65.6% |
| Female | 21 / 34.4% |
| **Race** |  |
| Black | 25 / 41% |
| White | 35 / 57.4% |
| Other | 1 / 1.6% |
| **Hemiparetic Side** |  |
| Right | 35 / 57.4% |
| Left | 26 / 42.6% |
| **\* Chronicity**<br>**(months post-stroke)** | 48.1 (46.9) [6-243] |
| **Lower Extremity**<br>**Fugl-Meyer Score** | 25.3 (6) [9-34] |
| **+ Self-Selected Overground**<br>**Walking Speed (m/s)** | 0.79 (0.29) [0.20-1.49] |

Continuous variables are presented as mean (standard deviation) [range]
Categorical variables are presented as frequency count / percentage
\* Missing date of stroke for 17 participants. All participants verbally confirmed their stroke occurred 6-months prior to enrollment.
+ Missing data for one participant

All participants provided written informed consent approved by the institutional review board at the Medical University of South Carolina. All individuals completed a biomechanical gait assessment without use of an AFO, clinical assessments including the lower extremity Fugl-Meyer (FMA-LE), Dynamic Gait Index (DGI), self-selected overground walking speeds (SS), and six-minute walk test (SMWT). Test-retest reliability of AAV and AAA were calculated from 23 stroke survivors who completed the biomechanical gait assessment on two separate days. Demographic characteristics are reported in Table 2.

**Table 2:** Participant Demographics in Test-Retest Analysis

| **n=23** |  |
| --- | --- |
| **Age** | 54.4 (14.8) [20-79] |
| **Sex** |  |
| Male | 14 / 60.9% |
| Female | 9 / 39.1% |
| **Race** |  |
| Black | 9 / 39.1% |
| White | 14 / 60.9% |
| **Hemiparetic Side** |  |
| Right | 18 / 78.3% |
| Left | 5 / 21.7% |
| **Chronicity**<br>**(months post-stroke)** | 52.9 (35.6) [8-135] |
| **Lower Extremity**<br>**Fugl-Meyer Score** | 23.9 (4.5) [15-33] |
| **Self-Selected Overground**<br>**Walking Speed (m/s)** | 0.87 (0.28) [0.42-1.27] |

Continuous variables are presented as mean (standard deviation) [range]
Categorical variables are presented as frequency count / percentage

Biomechanical data acquisition and analysis: Stroke survivors completed three 30-second trials of walking on a split-belt instrumented treadmill (Bertec Instrumented treadmill; Columbus, OH) at their self-selected walking speeds. Prior to data collection, participants walked on the treadmill during an acclimatization session to determine their self-selected speeds. Bilateral ground reaction force (GRF) data were sampled at 2000 Hz. Kinematic data were collected by using a motion capture system (PhaseSpace; San Leandro, CA, USA) using a modified Helen Hayes marker set and sampled at 120 Hz. A vertical GRF threshold of 1% body weight was used for identifying initial contact and gait events. The gait cycle was divided into six regions (first double support, first half of ipsilateral single leg stance, second half of ipsilateral single leg stance, second double support, first half of ipsilateral swing, second half of ipsilateral swing). We used the static standing calibration and functional joint centers for a link-based model. Foot clearance, ankle joint angles, angular velocity, and angular acceleration were then computed using a custom LabVIEW script (LabVIEW, National Instruments Corporation, Austin, TX). The peak foot clearance (FCL_P_), peak dorsiflexion angle (DFA_P_), peak AAV_P_, and peak AAA_P_ during the first half of ipsilateral swing of the paretic leg were calculated for each stride and averaged across strides and trials.

Measurement of Dorsiflexor Impairment: We used the FMA-LE to define dorsiflexor impairment with respect to the stepwise motor recovery from no dorsiflexion to the restoration of full dorsiflexor control. We calculated a measure of dorsiflexor impairment (FM_Rasch_) in Winsteps version 5.3.4 (Winsteps.com, Portland, OR) for each participant from the FMA-LE ankle items using the Andrich Rating Scale model (RSM). The RSM is an item response theory model that generates linear and interval measures from polytomous rating-scales and is a direct extension of the Rasch measurement model. The FMA-LE has previously been shown to fit the RSM for persons with chronic stroke [21]. Dorsiflexor impairment measures from the RSM were transformed from logits to traditional LE-FMA scores with 0 equating to no dorsiflexion control and 6 full dorsiflexor control [22].

Statistical Analysis: We used linear regression to quantify the amount of variance explained in dorsiflexion function (FM_Rasch_) by each biomechanical measure (independent variables; AAV_P_, AAA_P_, FCL_P_, DFA_P_). We also examined the amount of variance explained by speed (SS) to control for the speed dependent relationship of AAV_P_ and AAA_P_. We explored models controlling for speed to evaluate if speed alone explained statistically significant variance in FM_Rasch_. Statistical level of significance was set at p < 0.05. A forward stepwise regression was used to identify the model that captured the most variance in dorsiflexion function set at p<0.05. Pearson’s correlation was used to identify the relationship between clinical outcome measures and biomechanical measures. Strength of the correlations were defined as strong for coefficients ≥0.7, moderate for coefficients 0.4 < X <0.7, and weak for coefficients <0.3 [23]. William’s *t*-test [24] was used to statistically compare the magnitude of difference between two non-independent correlations with an overlapping variable. The correlation estimates between clinical outcome measures with AAV_P_ and AAA_P_ were compared to the correlation estimates found with FCL_P_ and DFA_P_.

To evaluate the test-retest reliability of AAV_P_ and AAA_P_ absolute agreement between repeated measures was tested with two-way mixed effects model for intraclass correlation coefficient (ICC) analysis.

All statistics were performed in SPSS version 25 (IBM Co., Somers, NY) and SAS version 9.4 (SAS Institute Inc., Cary, NC).

## Results

Regression analysis: We found that all models fit the assumptions for linear regression. AAV_P_ (Adj_R^2^ = 0.24; *p* < 0.0001), AAA_P_ (Adj_R^2^ = 0.2; *p* = 0.0002), FCL_P_ (Adj_R^2^ = 0.05; *p* = 0.0435), and DFA_P_ (Adj_R^2^ = 0.26; *p* < 0.0001) had a significant relationship with FM_Rasch_ while AAV_P_ and

DFA_P_ accounted for the most variance. We found a significant relationship between SS and FM_Rasch_ (Adj_R^2^ = 0.09; *p* = 0.0093). Stepwise regression revealed that speed was no longer significant and did not account for any variance when added to the model with AAV_P_ and DFA_P_.

Pearson’s correlation: We found that AAV_P_ and AAA_P_ had stronger correlation with most clinical outcome measures in comparison to FCL_P_ or DFA_P_. We found that FCL_P_ had a strong positive correlation with self-selected walking speeds (r = 0.71; *p* < 0.001) (Fig 1), moderate correlation with DGI (r = 0.46; p < 0.001) (Fig 4), SMWT (r = 0.49; p < 0.001) (Fig 3), and a weak correlation with FMA-LE (r = 0.37; *p* = 0.004) (Fig 2). DFA_P_ had a moderate positive correlation with FMA-LE (r = 0.42; *p* < 0.001) (Fig 2). A moderate positive correlation was also seen between AAV_P_ and DGI (r = 0.51; *p* < 0.001) (Fig 4), FMA-LE (r = 0.55; *p* < 0.001) (Fig 2), SMWT (r = 0.54; *p* < 0.001) (Fig 3), and self-selected walking speeds (r = 0.67; *p* < 0.001) (Fig 1). Lastly, we also found a moderately significant positive correlation between AAA_P_ and DGI (r = 0.56; *p* < 0.001) (Fig 4), FMA-LE (r = 0.56; *p* < 0.001) (Fig 2), SMWT (r = 0.61; *p* < 0.001) (Fig 3), and a strong correlation with self-selected walking speeds (r = 0.82; *p* < 0.001) (Fig 1).

**Figure 1.**
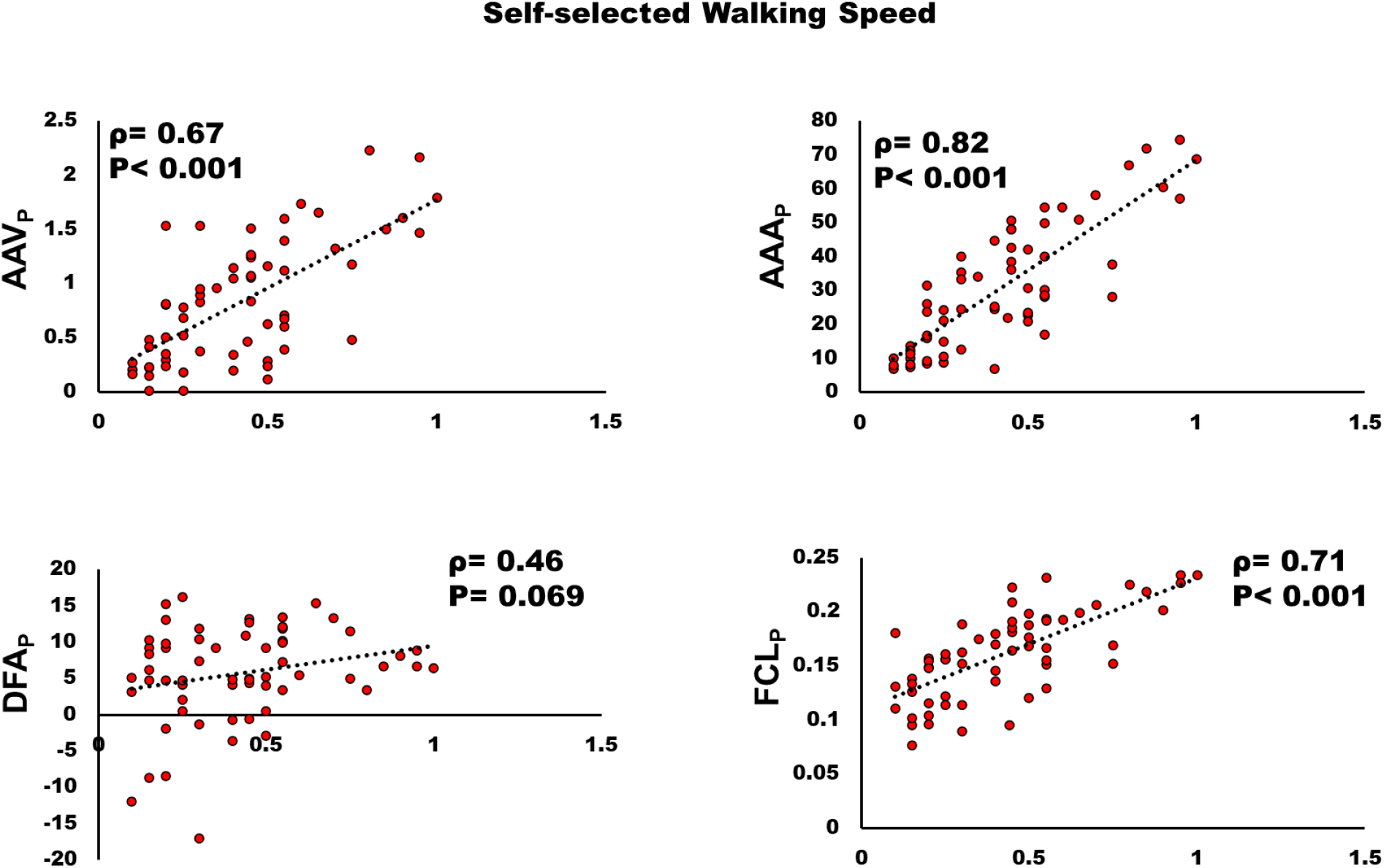
Relationship between self-selected overground walking speeds (SS) and A) peak ankle angular velocity (AAV_P_); B) peak ankle angular acceleration (AAA_P_); C) peak dorsiflexion angle (DFA_P_); D) peak foot clearance (FCL_P_). AAV_P_, AAA_P_, and FCL_P_ were positively correlated with SS (p<0.05), but no significant correlation was found between DFA_P_ and SS.

**Figure 2.**
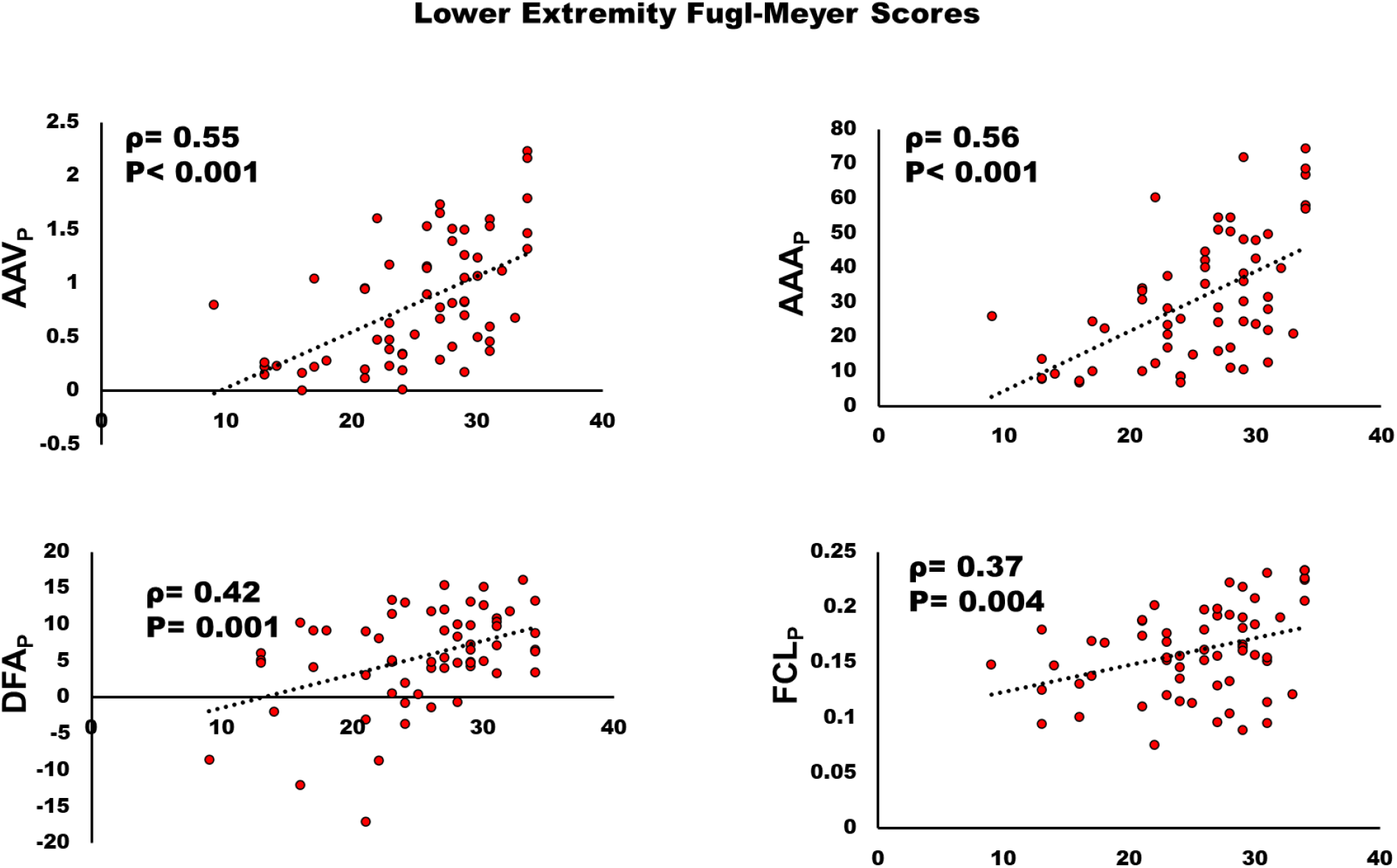
Relationship between lower extremity Fugl-Meyer (FMA-LE) and A) peak ankle angular velocity (AAV_P_); B) peak ankle angular acceleration (AAA_P_); C) peak dorsiflexion angle (DFA_P_); D) peak foot clearance (FCL_P_). All biomechanical measures were positively correlated with FMA-LE (p<0.05).

**Figure 3.**
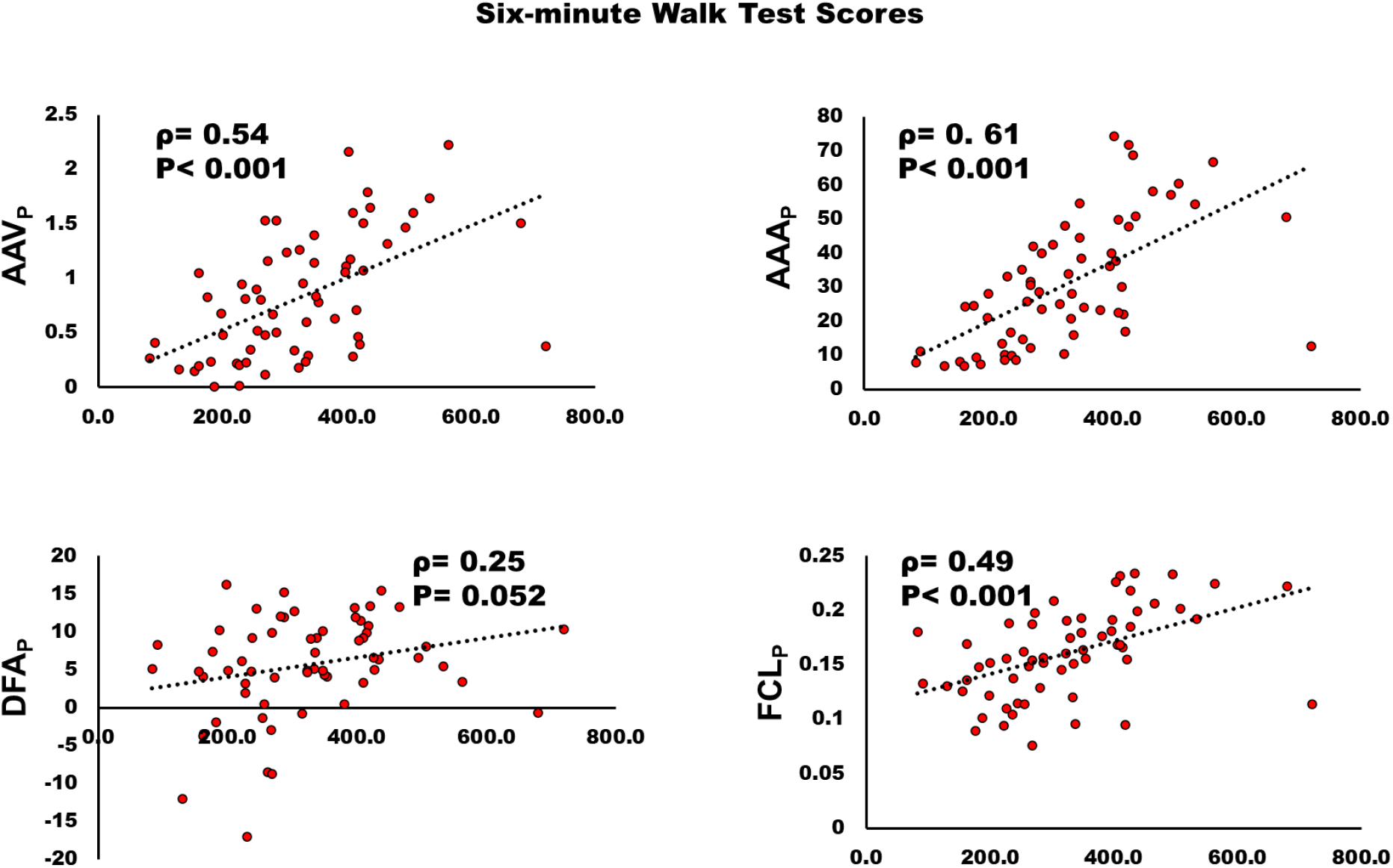
Relationship between six-minute walk test (SMWT) and A) peak ankle angular velocity (AAV_P_); B) peak ankle angular acceleration (AAA_P_); C) peak dorsiflexion angle (DFA_P_); D) peak foot clearance (FCL_P_). AAV_P_, AAA_P_, and FCL_P_ were positively correlated with SMWT (p<0.05), but no significant correlation was found between DFA_P_ and SMWT.

**Figure 4.**
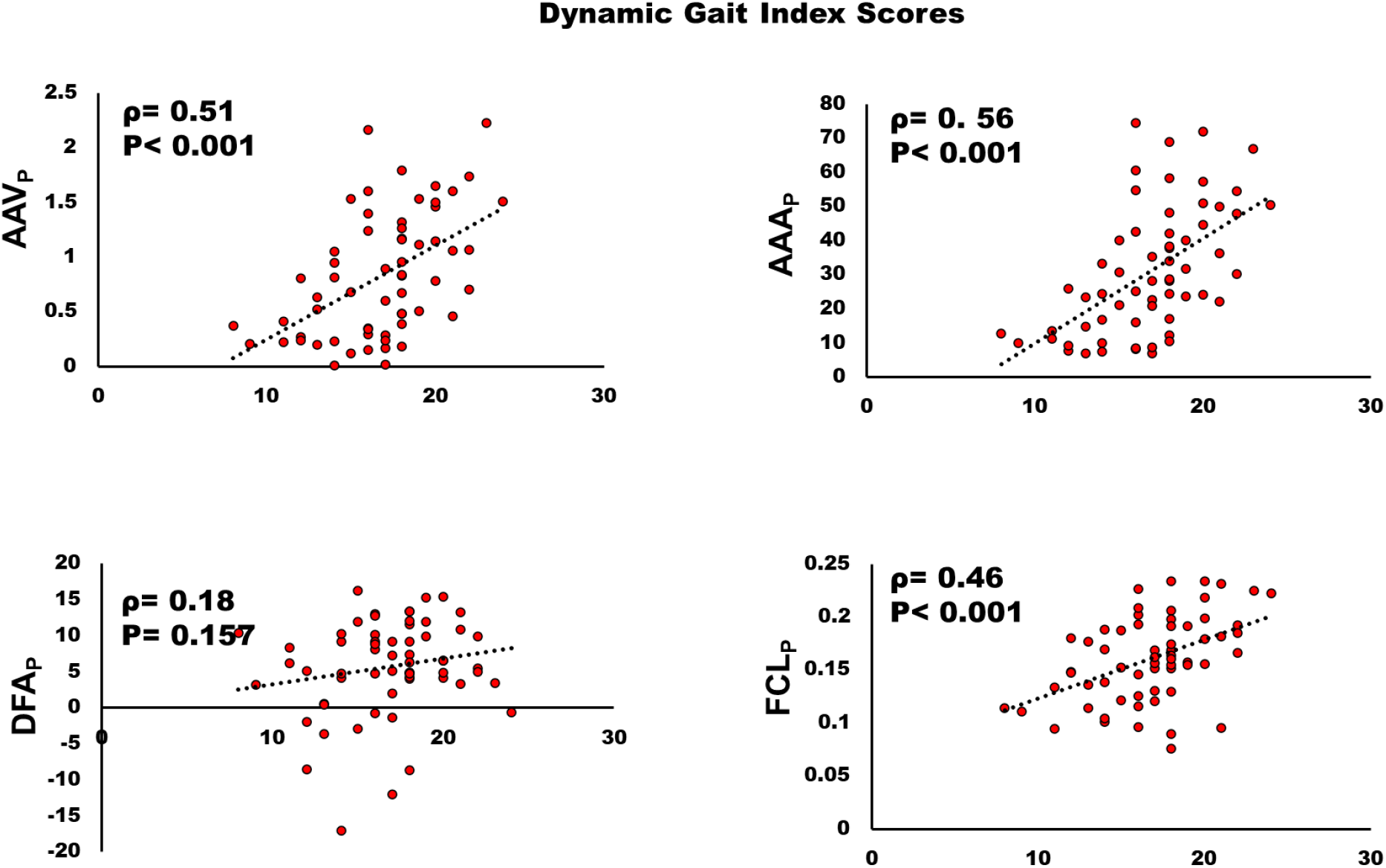
Relationship between Dynamic Gait Index (DGI) and A) peak ankle angular velocity (AAV_P_); B) peak ankle angular acceleration (AAA_P_); C) peak dorsiflexion angle (DFA_P_); D) peak foot clearance (FCL_P_). AAV_P_, AAA_P_, and FCL_P_ were positively correlated with DGI (p<0.05), but no significant correlation was found between DFA_P_ and DGI.

Post-hoc analysis with William’s *t*-test: The correlation coefficients for AAV_P_ and AAA_P_ were higher than FCL_P_ or DFA_P_ for most of the clinical outcome measures. Therefore, we used William’s *t*-test to conduct a statistical comparison of the magnitude of correlations between all clinical outcome measures with AAA_P_ and AAV_P_ compared to FCL_P_ and DFA_P_. We found that correlation between speed and AAV_P_ was significantly stronger than the correlation between speed and DFA_P_ by a magnitude of 0.44 (95% CI 0.18, 0.71; t_58_ = 3.42, p=0.001). The correlation between speed and AAA_P_ was significantly stronger than FCL_P_ by a magnitude of 0.11 (95% CI 0.02, 0.24; t_58_ = 2.54, p=0.01) and DFA_P_ by a magnitude of 0.59 (95% CI 0.35, 0.84; t_58_ = 5.58, p<0.0001). The correlation of DGI with AAV_P_ was significantly stronger than with DFA_P_ by a magnitude of 0.33 (95% CI 0.03, 0.61; t_58_ = 2.23, p=0.03), and with AAA_P_ was significantly stronger than with DFA_P_ by a magnitude of 0.37 (95% CI 0.08, 0.65; t_58_ = 2.60, p=0.01). We also found that the correlation of SMWT with AAV_P_ is stronger than with DFA_P_ by a magnitude of 0.29 (95% CI 0.01, 0.57; t_58_ = 2.07, p=0.04), and with AAA_P_ is stronger than with DFA_P_ by a magnitude of 0.36 (95% CI 0.08, 0.63; t_58_ = 2.63, p=0.01). The correlation of FMA-LE with AAV_P_ is stronger than with FCL_P_ by a magnitude of 0.19 (95% CI 0.03, 0.36; t_58_ = 2.34, p=0.02), and with AAA_P_ is stronger than with FCL_P_ by a magnitude of 0.20 (95% CI 0.06, 0.35; t_58_ = 3.04, p=0.004).

Test-retest reliability: The reliability analysis revealed excellent test-retest reliability for both AAV_P_ (ICC = 0.968) and AAA_P_ (ICC = 0.947) calculated across gait cycles during the 30 seconds trials.

## Discussion

In the current study, we investigated whether AAV_P_ and AAA_P_, could be used as a measure for dorsiflexion function during walking. We compared the performance of AAV_P_ and AAA_P_ with FCL_P_, and DFA_P_ in explaining dorsiflexion function and their association with clinical measures of walking ability. Our results revealed that AAV_P_, AAA_P_, FCL_P_, and DFA_P_ were all significantly related to impaired dorsiflexion function evaluated using FM_Rasch_. Stepwise linear regression revealed that AAV_P_ and DFA_P_ explained approximately half the variance for FM_Rasch_. We also found that speed was not a significant co-variate in the model suggesting that speed dependency is not influencing our findings. Our correlation analysis showed that both AAV_P_ and AAA_P_, had stronger correlations with post-stroke walking ability measures compared to FCL_P_ and DFA_P_. Lastly, we found that AAV_P_ and AAA_P_ had excellent test-retest reliability. Together this suggests that AAV_P_ and DFA_P_ may best represent dorsiflexion impairment with respect to the FMA-LE based stepwise motor recovery i.e., FM_Rasch_ in individuals with stroke. Additionally, AAV_P_ and AAA_P_ represent walking ability better than FCL_P_ and DFA_P_.

Our findings indicate that DFA_P_ and FCL_P_ may not be ideal outcome measures to evaluate the effects of impaired dorsiflexion function during swing on walking ability. Although, DFA_P_ and FCL_P_ were significantly related to impaired dorsiflexion function, they both had weak to moderate correlations with clinical measures of walking ability. Kesar et al., [25] have shown increases in peak dorsiflexion angle following FES during walking, but they did not report any changes that translate into an improvement in quality of life. Our results show DFA_P_ has weak or no correlation with functional walking ability measure. Therefore, it is possible that DFA_P_ predicts impaired dorsiflexion function measured as FM_Rasch_ which is based on FMA-LE ankle dorsiflexion function, but it was not significantly associated with walking ability. We also found that FCL_P_ performed better than DFA_P_ when correlating with walking ability measures. However, FCL_P_ does not solely represent the dorsiflexor function, but is associated with contribution from hip, knee, and ankle joint movements during swing [26]. Therefore, FCL_P_ and DFA_P_ may not be as directly associated with dorsiflexor function during walking as AAV_P_ and AAA_P_.

Both AAV_P_ and AAA_P_ have the potential to be used in the future as clinical outcome measures of dorsiflexion function during walking in the stroke population. In the current study both AAV_P_ and AAA_P_ were computed based on the 3D kinematic and kinetic data, but there is a potential for AAA_P_ to be computed using tri-axial accelerometers in a clinical set-up. AAA_P_ as measured using accelerometers could be a potential outcome measure that would give clinicians an opportunity to evaluate dorsiflexion function during walking to optimize individualized treatment protocol and modulate it with progression of treatment as needed.

In conclusion, AAV_P_ and AAA_P_ can be used as measures of impaired dorsiflexion function during walking in the stroke population. However, there are some limitations to be considered. Moderate and stronger correlations of walking speeds with AAV_P_ (r = 0.67; *p* < 0.001) and AAA_P_ (r = 0.82; *p* < 0.001) respectively should be interpreted with caution as both are speed dependent variables. However, our regression models revealed that speed explained only 9% of variance in FM_Rasch_ (Adj_R^2^ = 0.09; *p* = 0.0093) in comparison to 24% explained by AAV_P_ (Adj_R^2^ = 0.24; *p* < 0.0001), and 20% explained by AAA_P_ (Adj_R^2^ = 0.2; *p* = 0.0002).

Additionally, speed did not account for any variance when added to the stepwise regression model with AAV_P_ and DFA_P_. This implies that AAV_P_ and AAA_P_ measures can be interpreted regardless of speed for quantifying dorsiflexor function with respect to FM_Rasch_. Another consideration when interpreting current results is that there is no gold standard outcome measure to evaluate the impaired dorsiflexion function during walking which limits our ability to test the validity of AAV_P_ and AAA_P_. However, results from the current study will provide researchers and clinicians with reliable measures that can evaluate the effects of impaired dorsiflexion function on walking ability with the potential of AAA_P_ to be used in a clinical set-up. Future studies are needed to validate AAA_P_ recorded from accelerometers against AAA_P_ computed with 3D based kinematic and kinetic data captured with the motion capture system. In addition to being easy to set-up in the clinic, accelerometers have the potential to be incorporated with portable stimulation devices that can be used for advanced, feedback-controlled FES applications [27]. Furthermore, with accelerometers the performance of dorsiflexion function can also be assessed on inclined planes, uneven terrains, stairs etc. which will be more relevant to the daily activities of stroke survivors.

## Disclaimers

Any opinions expressed in this work are those of the authors and do not necessarily reflect the view of the U.S. Department of Veteran Affairs, or the NIH.

## Authors’ contributions

SS, CC, JK, SK, and MB have developed the concept and design of the study. SS, AB, BS, JK, and MB have analyzed the data and SS has drafted the manuscript. SS, BS, SK, CC, and MB have been involved with designing the study protocol and acquisition of data. All authors have revised the manuscript critically to improve intellectual content and approved the final manuscript.

## Acknowledgements

The authors would like to thank Heather Knight at the Medical University of South Carolina for helping with the data analysis.

